# Flexible docking of cyclic peptides to proteins using CABS-dock

**DOI:** 10.1101/2025.06.18.660300

**Authors:** Mateusz Zalewski, Aleksandra Badaczewska-Dawid, Sebastian Kmiecik

## Abstract

Cyclic peptides are promising therapeutics, but their flexible docking remains challenging. We present a protocol based on the well-established CABS-dock method, enhanced with cyclic restraints and Rosetta refinement. The approach was evaluated on 38 benchmark complexes previously used in other docking method studies. While selecting the truly best model remains difficult, near-native solutions are frequently sampled. CABS-dock offers global, unbiased docking without prior binding site knowledge, making it valuable for pose generation, structural ensemble modeling, and integration into AI-driven peptide–protein docking workflows.

## Introduction

Peptides are highly promising drug candidates due to their strong target specificity, low toxicity, and compatibility with rational design strategies^1^. However, linear peptides often suffer from major pharmacological limitations, including conformational flexibility, proteolytic instability, and poor oral bioavailability and membrane permeability^2,3^. Cyclization - through backbone or disulfide linkages - enhances peptide rigidity, stability, and target affinity^4^, making cyclic peptides promising tools for modulating protein–protein interactions.

Among the early computational tools addressing this challenge, AutoDock CrankPep (ADCP) was one of the first capable of handling flexible cyclic peptides, and it remained the state-of-the-art for several years^5,6^. More recently, the field has shifted toward deep learning–based approaches, for both linear^7,8^ and cyclic peptides^9,10^. For cyclic peptides, HighFold^9^ has shown consistently higher accuracy than earlier methods, including ADCP^5^ and AfCycDesign^10^ (an AlphaFold based protocol), in benchmarks of cyclic protein-peptide complexes. In particular, it achieved superior Top-1 success rates and interface-level accuracy, establishing a new performance baseline in this domain.

Despite this progress, AI-based methods like HighFold are typically deterministic, producing a single binding pose per input. This can be limiting in cases where binding is ambiguous or flexible, or where pose diversity and interpretability are crucial. In contrast, methods, though generally inferior in scoring accuracy, can offer extensive conformational sampling that captures a broader landscape of potential binding modes.

In this work, we present a flexible docking protocol for cyclic peptide–protein complexes that supports both local and global docking modes. The global setup, which does not rely on prior knowledge of the binding site, represents - to our knowledge - the first such application for cyclic peptides. The protocol builds upon the well-established CABS-dock framework□^11–13^ for flexible protein–peptide docking based on coarse-grained modeling and Monte Carlo sampling. To accommodate cyclic topologies, we extended CABS-dock with (1) distance restraints to preserve ring closure during sampling, (2) PD2-based reconstruction of Cα traces to all-atom resolution^14^; and (3) Rosetta FlexPepDock-based refinement for improved backbone and side-chain accuracy^15,16^. We evaluated both docking modes on a benchmark of 38 experimentally determined cyclic peptide–protein complexes from the ADCP dataset^5^, 17 of which overlap with the HighFold study^9^.

## Materials and methods

CABS-dock^11–13^ is a multiscale molecular docking method that employs a coarse-grained simulation engine based on the CABS model, illustrated in Supplementary Figure 1, and uses Replica Exchange Monte Carlo sampling to explore peptide–protein interactions. Within the method, the resulting models are reconstructed to all-atom resolution. Originally developed as a web server for global, flexible docking^11,12^, it has since been adapted to a range of modeling tasks, including peptide docking with structural information^17^, protein–protein docking^18,19^, GPCR–peptide interactions^20^, docking involving large receptor conformational changes^21^, and peptide cleavage site prediction^22^. A standalone version^13^ allows for full control over simulation parameters and custom workflows.

To support cyclic peptides, we introduced distance restraints to maintain the closed-ring topology of backbone-cyclized and disulfide-cyclized peptides. These restraints were defined using the built-in constraint system of the standalone version (See Supplementary Methods for detailed implementation and example syntax).

The simulation involves multiple steps, including conformational sampling, clustering, and all-atom reconstruction (see Supplementary Figure 2). Each docking simulation generated 10,000 Cα-trace models, sampled without prior knowledge of the binding site. The models were scored using CABS internal energy terms, followed by structural clustering to identify the top 10 candidates.

In this work, we applied two docking modes: global and local. In global docking, the peptide was allowed to interact with the entire receptor surface. In local docking, the peptide was spatially restrained to the binding site region but remained fully flexible during sampling. This setup follows a protocol previously applied to linear peptides^17^ (see Supplementary Methods for implementation details). Figure□1 presents an example case from the benchmark, highlighting the spatial extent of peptide sampling in global versus local docking setups.

**Figure 1.**
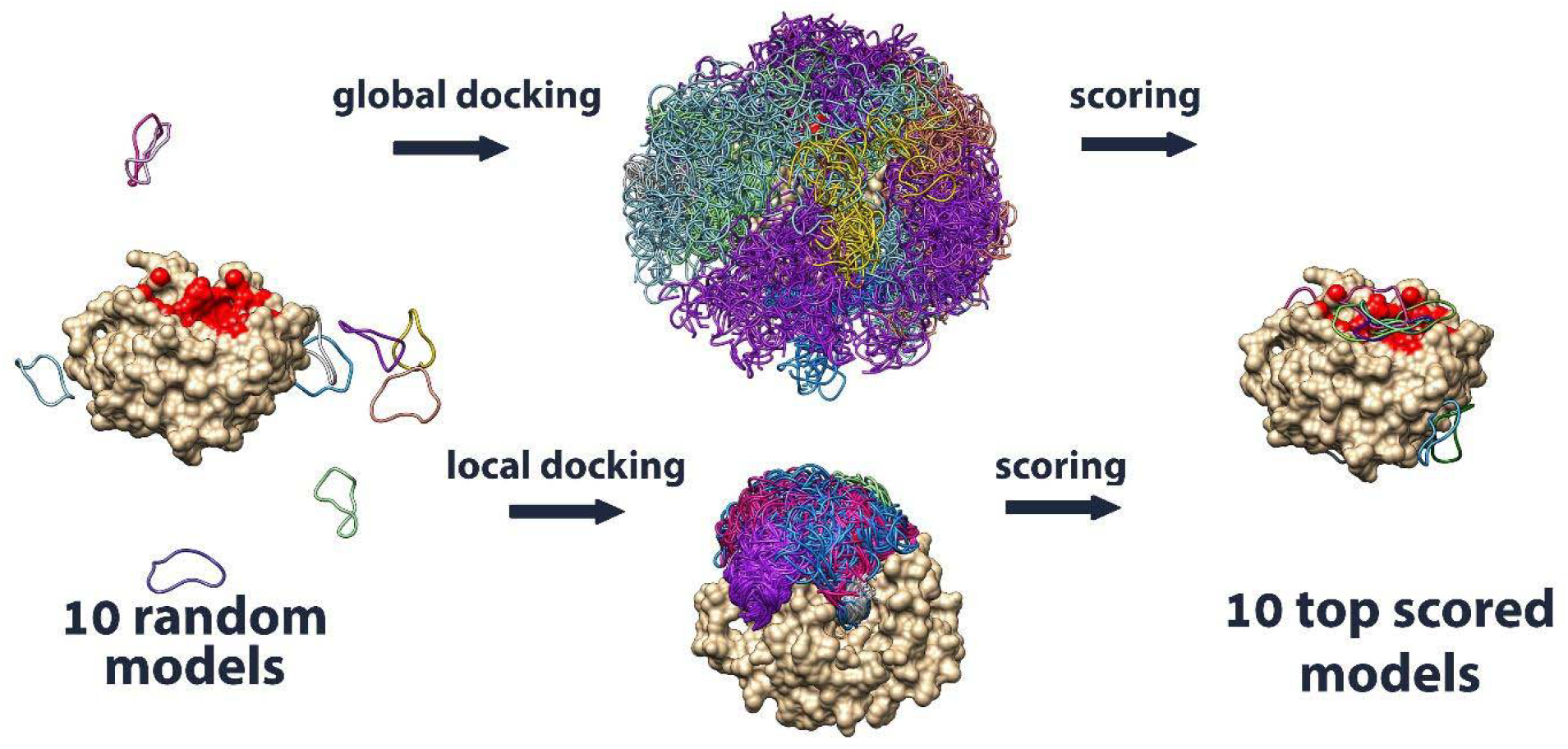
Visualization of peptide sampling in global and local docking modes, shown for a representative benchmark complex.

### Benchmark set and evaluation

We evaluated the method on 38 peptide–protein complexes from the ADCP benchmark set^5^, comprising 18 backbone-cyclized and 20 disulfide-cyclized peptides. Each complex was redocked to both the bound (*holo*) and unbound (*apo*) receptor conformations. Success was measured using the CAPRI-defined Fnat metric^23^, with additional ensemble analysis based on LRMSD (details in SI).

## Results and discussion

In this work, we evaluated flexible docking of cyclic peptides to both *holo* and *apo* receptor structures using CABS-dock, in global and local modes. To our knowledge, this represents the first application of global docking to cyclic peptides without prior knowledge of the binding site. The benchmark included 38 complexes, divided into backbone-cyclized (18) and disulfide-cyclized (20) cases. Apo forms were available for 17 backbone-cyclized and 10 disulfide-cyclized complexes^5^. For each target, docking simulations were run in both modes, and the top 10 models were reconstructed from Cα traces to all-atom resolution and refined with Rosetta FlexPepDock. This refinement step was previously applied to linear peptides in multistage protocols for GPCR–peptide docking^20^ (see Supporting Information for details). In our benchmark, the refined models reached acceptable quality (Fnat□≥□0.3) in 83% of backbone-cyclized and 80% of disulfide-cyclized *holo* complexes, with 67% and 35% reaching medium or high accuracy (Fnat□≥□0.5). For *apo* structures, the corresponding rates were 82% and 70%, with 56% and 50% reaching Fnat□≥□0.5. These values reflect the best-performing models selected from the top 10 scoring models in each simulation. Full results are provided in Supplementary Tables□1 and□2.

To complement the analysis of top-scored models, we evaluated the quality of the full CABS-dock output ensemble (top 10,000 models per run) using LRMSD, which is more suitable for C-alpha traces than Fnat, as it does not rely on atom-level contact definitions and provides a reliable measure of backbone-level structural accuracy. Average LRMSD values on bound complexes were 3.89 Å (cyclized by backbone) and 5.09 Å (cyclized by disulfide bond) for global docking, and 2.65 Å (backbone) and 4.23 Å (disulfide) for local docking. On unbound receptors, the respective averages were 4.76 Å (backbone) and 6.46 Å (disulfide) for global docking, and 2.92 Å (backbone) and 3.82 Å (disulfide) for local docking. These values confirm that even when top-scoring models miss the native state, the full ensemble often includes high-quality candidates (see Figure 2A).

**Figure 2.**
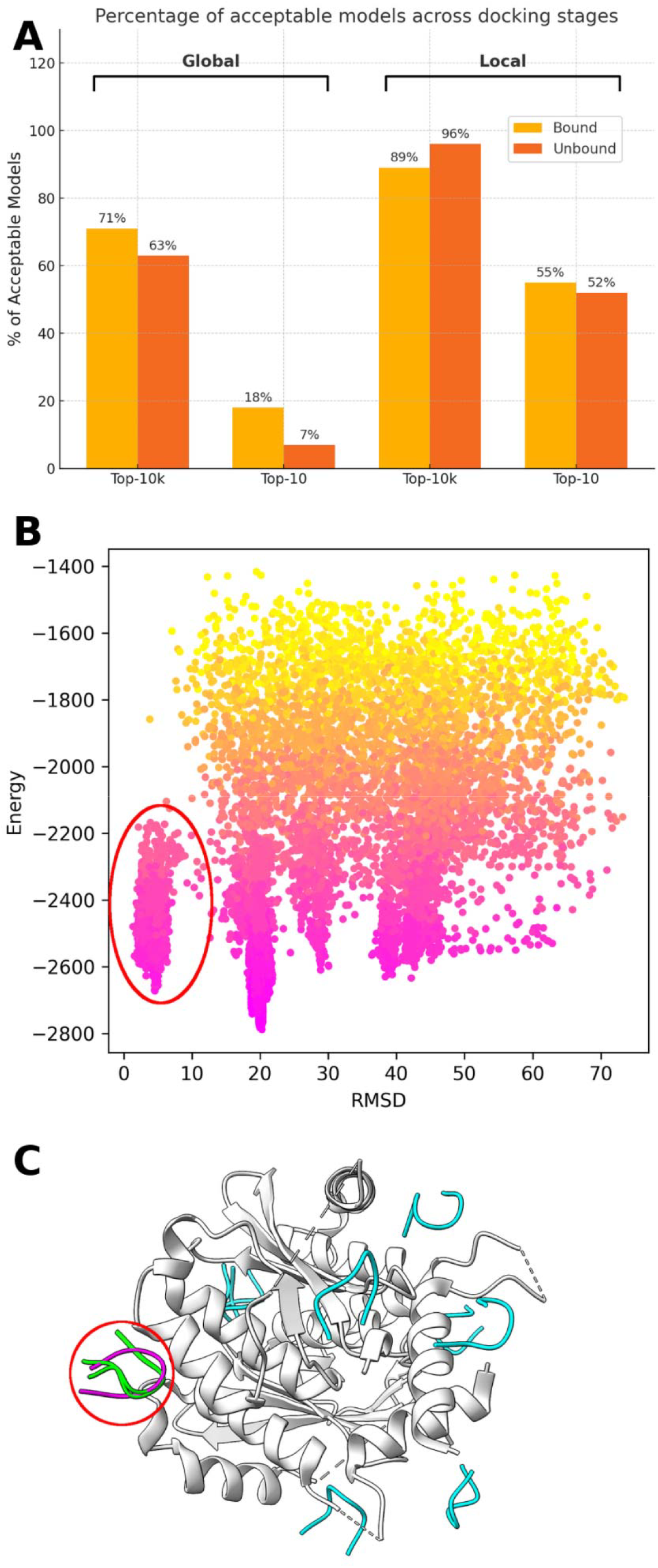
Sampling and scoring behavior in CABS-dock. **A:** Fraction of complexes with at least one acceptable model (LRMSD <□5.5□Å) in Top-10 vs Top-10k, for global and local docking, and bound/unbound receptors. **B:** Energy vs. LRMSD for complex 3WNE (global docking, bound receptor). Near-native models are circled in red. **C:** Molecular visualization of the same case as in B. Top-10 predicted models are shown, with near-native structures in green and others in cyan. Experimental structure shown in magenta; red circles indicate near-native models.

Using the 5.5 Å LRMSD threshold established by Raveh et al.^24^ as a reliable basin of attraction for near-native refinement, we found that for bound cases, acceptable structures (<5.5 Å) were present in 27 out of 38 cases (71%) with global docking, and in 34 out of 38 cases (89%) with local docking. For unbound cases, these numbers were 17 out of 27 (63%) for global and 26 out of 27 (96%) for local docking.

In summary, while near-native models are frequently generated, they are not always selected as top-ranked due to limitations in the scoring procedure. In CABS-dock, scoring is based on structural clustering combined with energy evaluation in the coarse-grained CABS model. Similar scoring-related issues - where accurate models were generated but not top-ranked - have also been observed previously for docking of linear peptides^11,12,17^. Figure□2A shows that more accurate models often exist among the 10,000 generated structures than among the top 10 selected ones, indicating that the main limitation lies in scoring rather than sampling. Panels□B and□C illustrate a representative case where a near-native model appears in the top 10 but is not ranked first. Instead, the top-scoring structure is a misdocked conformation, likely favored due to its larger population and a deeper minimum in the coarse-grained energy landscape.

### Comparison with ADCP

To assess the performance of CABS-dock relative to established docking tools, we compared it directly with AutoDock CrankPep (ADCP) - the previous state-of-the-art method for flexible docking of cyclic peptides prior to the emergence of AI-based predictors. The comparison was performed on the full benchmark set of bound (*holo*) and unbound (*apo*) receptor structures, grouped by peptide cyclization type: backbone or disulfide. For both methods, results are reported for the best model among the top 10 scored predictions, following standard procedures from the original ADCP study^5^ □

For bound complexes, CABS-dock showed competitive performance. In the backbone-cyclized subset (n□=□18), it achieved an average Fnat of 0.61, with 83% acceptable (Fnat□≥□0.3) and 67% medium-quality (Fnat□≥□0.5) predictions. ADCP performed slightly better overall (avg Fnat 0.72; 100% acceptable; 78% medium), but CABS-dock outperformed ADCP in 7 out of 18 cases, indicating comparable strength in certain structural contexts. In the disulfide-cyclized subset (n□=□20), CABS-dock yielded an average Fnat of 0.43 (16 acceptable, 7 medium), versus 0.60 for ADCP (19 acceptable, 15 medium). Again, CABS-dock surpassed ADCP in 7 targets, reinforcing its utility across structurally diverse or flexible systems.

For unbound complexes, both methods showed reduced performance due to increased receptor flexibility. The drop was more pronounced for CABS-dock, particularly in scoring. In the backbone-cyclized unbound set (n□=□17), CABS-dock reached an average Fnat of 0.53 (14 acceptable, 10 medium), while ADCP maintained higher accuracy (avg Fnat 0.70; 17 acceptable; 15 medium). CABS-dock performed better on 4 targets. In the disulfide-cyclized unbound group (n□=□10), CABS-dock averaged 0.35 (7 acceptable, 2 medium), compared to ADCP’s 0.49 (10 acceptable, 4 medium), with CABS-dock outperforming ADCP in 3 cases.

While average performance favors ADCP, CABS-dock consistently generates near-native models, especially in challenging unbound scenarios, thanks to its flexible Monte Carlo sampling. However, limitations in scoring and model selection often prevent these models from being top-ranked. This is further supported by Figure□2A, where a greater number of unbound cases contain acceptable models among the Top-10 predictions compared to the bound set. Importantly, average performance does not capture the full picture of target-specific variability.

As shown in Figure□3, per-target Fnat values vary widely across both methods, with each outperforming the other in a subset of cases. This suggests that their complementary strengths could be leveraged through selective or combined use to improve overall reliability.

**Figure 3.**
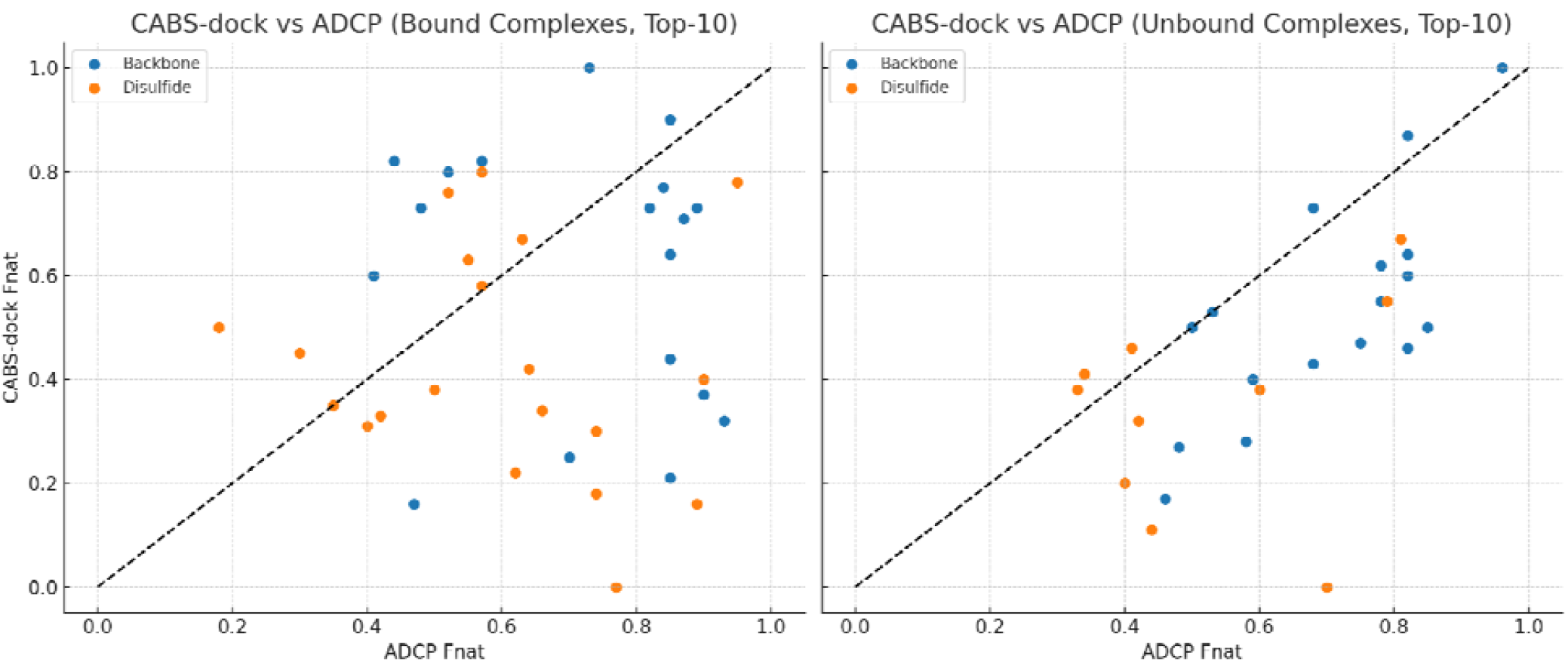
Per-target comparison of CABS-dock and ADCP (Top-10 Fnat) on bound (left) and unbound (right) complexes. Each point represents one target; colors indicate peptide cyclization type. Dashed line marks equal performance.

## Comparison with HighFold

To evaluate the performance of CABS-dock against modern AI-based methods, we compared it to HighFold - a recent deep learning framework for docking cyclic peptides. HighFold has shown superior predictive accuracy over previous methods, including AlphaFold-based AfCycDesign method^10^. We used the same benchmark set of 17 unbound backbone-cyclized protein–peptide complexes and evaluated only the top-ranked (Top-1) models from each method, as reported in the HighFold study^9^. Acceptable models were defined using an Fnat threshold of 0.2, following the HighFold benchmark. This differs from the 0.3 threshold used in earlier comparisons to ADCP, which followed its original evaluation protocol^5^.

HighFold achieved acceptable predictions (Fnat ≥ 0.2) in all 17 cases (100%), with an average Fnat of 0.82, indicating consistently high accuracy. CABS-dock, by comparison, reached this threshold in only 7 cases (35.3%), with an average Fnat of 0.24. This gap reflects core methodological differences: HighFold directly predicts refined backbone structures using a transformer model trained on cyclic peptide complexes, while CABS-dock relies on exhaustive sampling with a coarse-grained model, followed by reconstruction and refinement. Although CABS-dock generates 10,000 models per run, its scoring function often fails to identify the most accurate structures.

Nevertheless, in several cases, CABS-dock’s broad sampling successfully recovered viable binding modes. A notable example is the 3ZGC complex: both methods produced acceptable Top-1 models, though none reached the medium-quality threshold (Fnat ≥ 0.5). This was the worst-performing case in the HighFold benchmark, due to its unusual binding mode - an extended peptide conformation at the receptor rim - which is difficult for models trained primarily on pocket-like interactions^9^. HighFold, AlphaFold, and CABS-dock all achieved Fnat values around 0.44–0.47, while ADCP scored lower (0.15). Importantly, CABS-dock’s Top-10 ensemble included better-quality poses, reaching Fnat = 0.53 (unbound) and 0.85 (bound). These results show that while CABS-dock lags behind in Top-1 accuracy, it can still generate near-native solutions in challenging cases. Supplementary Figure 3 illustrates this contrast across all targets, with 3ZGC emerging as a unique outlier. Although HighFold excels in predictive precision, CABS-dock may contribute valuable structural diversity in hybrid or ensemble-based workflows.

## Conclusions

In this study, we present a proof-of-concept application of the CABS-dock method for docking flexible cyclic peptides to protein targets. The protocol employs coarse-grained Monte Carlo sampling with distance-based restraints to mimic cyclic topology, followed by all-atom reconstruction. This work builds directly upon our previous studies, in which we used the standalone CABS-flex engine to predict the structures of free linear and cyclic peptides^25^, as well as for docking linear peptides to globular proteins and GPCRs^20^. Here, we adapt the same core methodology to the docking context, demonstrating that flexible sampling can be applied to model bound-state peptide conformations under topological constraints.

Although the overall accuracy of predicted complexes does not match that of deep learning models such as AlphaFold or HighFold, CABS-dock performs comparably to classical methods such as ADCP, while offering a key advantage: the ability to explore diverse peptide conformations under cyclic constraints. This makes it particularly useful in exploratory modeling where binding modes are unknown, or when structural ensembles are needed for re-ranking, refinement, or experimental data interpretation. Unlike many AI-based structure prediction tools, which are deterministic and often opaque, CABS-dock offers full control over flexibility, restraints, and sampling settings. This configurability supports its use in integrative or hypothesis-driven modeling, especially when peptide flexibility is biologically relevant.

Experimental evidence supports the idea that conformational plasticity plays a key role in cyclic peptide function, permeability, and binding specificity. At the same time, predictors like AlphaFold often over-stabilize ordered structures and miss alternative or transient states observed by NMR or MD^26^ By explicitly sampling such conformational alternatives, CABS-dock has the potential to address this gap. While our evaluation focused on local docking near known interfaces, the method also supports global docking without prior knowledge of the binding site - making it applicable to challenging or non-canonical targets. Although its Top-1 ranking accuracy remains lower than that of state-of-the-art AI models, near-native solutions often appear in the generated ensembles, underscoring its value in hybrid or ensemble-based pipelines.

Looking ahead, further development could include guiding sampling with experimental restraints (e.g., from NMR or crosslinking) and improving model selection using physics-based or machine learning scoring. As such, CABS-dock should be seen not only as a docking tool, but also as a generator of diverse, physically plausible conformational states. This flexibility, combined with support for both local and global docking under cyclic constraints, gives it unique utility in exploratory modeling, data integration, and studies of systems where structural heterogeneity is functionally important.

## Supporting Information

Supplementary Information includes detailed descriptions of the docking workflow, input preparation, and reconstruction/refinement protocols used in this study. It also provides comprehensive tables reporting docking performance across all benchmark cases, as well as schematic figures illustrating the CABS model and simulation steps.

## Acknowledgments

Mateusz Zalewski and Sebastian Kmiecik have been supported by the National Science Centre, Poland (OPUS 2020/39/B/NZ2/01301).

## Data and Software Availability

Detailed instructions for reproducing the docking protocol are provided in the Supplementary Information (SI) of the manuscript. The standalone version of CABS-dock used in this study is available at: https://bitbucket.org/lcbio/cabsdock. All output structures corresponding to the Top-10 predicted models for each target are available in the associated GitHub repository: https://github.com/ZalewskiMa/CABSdock-cyclic.

