## Supplementary Information for "Flexible docking of cyclic peptides to proteins using CABS-dock"

**Supplementary Methods**

The CABS-dock method is built on the CABS model (C-alpha, C-beta, Side-chains)[1], which represents each amino acid using up to four pseudo-atoms (see Supplementary Figure 1). CABS-dock employs a knowledge-based statistical potential and an efficient Replica Exchange Monte Carlo (REMC) scheme for conformational sampling.

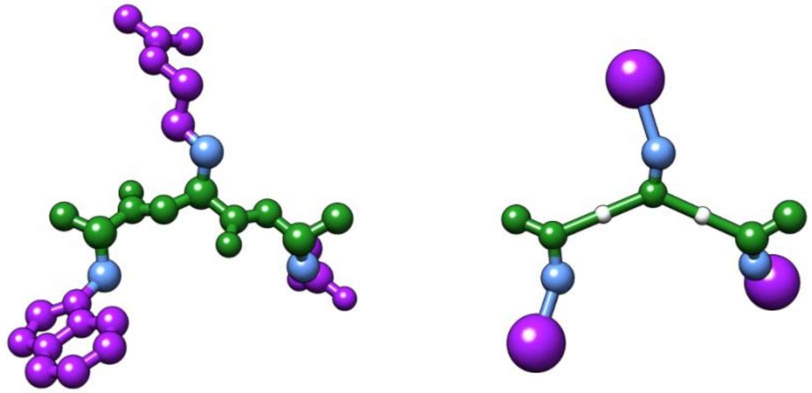

**Supplementary Figure 1.** Schematic illustration of protein chain fragment in all-atom (left) and CABS (right) model representation. In the CABS model, each residue is represented by up to four pseudoatoms: Cα (alpha carbon), Cβ (beta carbon), a pseudoatom for the side chain center, and a virtual center of the peptide bond.

**CABS-dock Simulation Workflow**

The CABS-dock simulation involves five main steps (Supplementary Figure 2):

1. **Setting starting conformations.** The algorithm generates ligands starting structures at random positions within about 20 Å from the receptor. Starting structures are represented by C-alpha traces.

2. **Conformational space sampling.** The docking simulation is performed using the REMC sampling scheme. Models generated during sampling are saved into a pseudo-trajectory for every starting structure.

3. **Reconstruction to CABS representation.** All generated during simulation models are reconstructed from C-alpha traces to CABS representation.

4. **Scoring.** CABS-dock scoring is a two-step function. During the first stage 100 conformations of protein-peptide complexes are selected, based on their CABS energy values (the lower the interaction energy the better) from every trajectory. The second stage is structural clustering using a k-medoid algorithm, leading to the 10 best structures selected.

5. **Reconstruction to all-atom representation.** Top-scored models are rebuilt to an all-atom representation. By default, reconstruction is performed using a Modeller rebuilding procedure[2]. Please note that in this work reconstruction was carried out using the PD2[3], followed by high-resolution refinement with Rosetta FlexPepDock[4], which optimizes backbone geometry and side-chain packing at the protein–peptide interface.

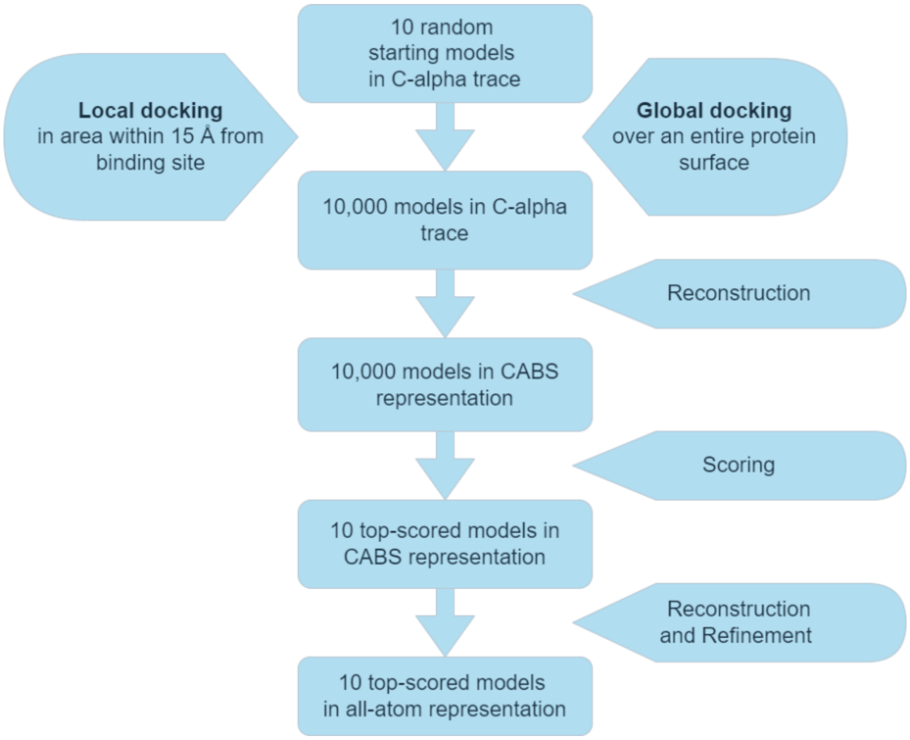


**Supplementary Figure 2.** Steps of molecular docking simulation using CABS-dock.

**Distance Restraints for Cyclization and Local Docking**

The CABS-dock tool has been designed for docking linear peptides, so we had to provide a solution to the problem of preserving the cyclic ring of peptides during simulation. It was achieved by using distance restraints offered by CABS-dock standalone. Standard command line for running CABS-dock simulation:

*CABSdock –i protein-pdb-code –p peptide-sequence:peptide-secondary-structure*

can be enriched with additional flag:

*--ca-rest-file FILE*

which allows adding distance restraints from a file. *FILE* should contain information about (*dist* ± *tolerance*) Å distance restraint between CA atom in residue *resi* and CA atom in *resj*, and restraint weight *weightm* for lesser distance or *weightn* for greater distance; in the following order:

*resi resj dist tolerance weightm weightn*

For instance, to add (3.78 ± 0.5) Å distance restraint between CA atoms of first and seventh amino acid in peptide and to set this restraint weight to 2, restraints-file should contain following text:

*1:PEP 7:PEP 3.78 0.5 2.0 2.0*

Default CABS-dock simulation allows peptides to interact with the entire receptor interface. However, it is possible to restrict conformational space by adding distance restraints. It can be done the same way as preserving the cyclic ring of peptides during simulation. Single restraints-file can contain information about multiple restraints, so there is no need to add multiple *--ca-rest-*

*file* flags and multiple files. In order to perform local docking restraints-file should contain strong restraints (*weightn* = 100) between residue resi of receptor and all peptide residues (*1:PEP*, *2:PEP*, *3:PEP*, …), which do not allow peptide to move away from residue *resi* further than *dist*. For instance, restraints-file for local docking head-to-tail peptide of 7 amino acids length to a receptor, within 15 Å from fiftieth amino acid of receptor's chain A, restraints-file should contain the following text:

*1:PEP 7:PEP 3.78 0.5 2.0 2.0*
*50:A 1:PEP 15 0.0 0.0 100.0*
*50:A 2:PEP 15 0.0 0.0 100.0*
*50:A 3:PEP 15 0.0 0.0 100.0*
*50:A 4:PEP 15 0.0 0.0 100.0*
*50:A 5:PEP 15 0.0 0.0 100.0*
*50:A 6:PEP 15 0.0 0.0 100.0*
*50:A 7:PEP 15 0.0 0.0 100.0*

**Global and Local Docking Setups**

In this work we have performed both global and local docking using the CABS-dock method. Restraints-file for global docking simulations contained only information about restraints responsible for peptide cyclization. Restraints-file for local docking simulations contained the same restraints as their equivalent for global docking and additional restraints responsible for keeping peptide structure in the desirable area (20 Å for peptides longer than 14 amino acids and 15 Å for all other peptides). Schematic visualization of cyclic peptides global and local docking is shown in Figure 1.

**Reconstruction and Refinement**

The top 10 Cα-trace models from each CABS-dock simulation were reconstructed and refined following the protocol described by Badaczewska-Dawid et al.[5]. Backbone reconstruction was carried out using PD2[3], which builds full backbone atom positions based on statistical modeling of local geometry using Gaussian mixture models. Side chains were subsequently added using SCWRL4, which selects the most favorable rotamers based on backbone-dependent rotamer libraries and steric optimization. The resulting all-atom models were further processed with Rosetta FlexPepDock[4], employing its high-resolution refinement protocol. This includes Monte Carlo-based backbone perturbation, side-chain repacking, and energy minimization of the entire interface using the Rosetta full-atom scoring function. Importantly, the peptide backbone was allowed to move flexibly during refinement, while the receptor was kept fixed.

**Dataset**

Proposed protocol for docking cyclic peptides using CABS-dock was validated using a benchmark described by Zhang et al.[6] Dataset consists of 38 cyclic peptide complexes, of which 18 are cyclized by their backbone and 20 by disulfide bond. All of the structures contained in the benchmark have crystallographic resolution values equal or above 2.2 Å and no alternative locations or missing atoms in their PDB entries; and all of the peptides consist of 5 to 20 standard amino acids.

**Quality measures**

Typically, evaluation of a docking success is performed using CAPRI (Critical Assessment of Predicted Interactions) criteria[7]: IRMSD (Interface Root Mean Square Deviation), LRMSD (Ligand Root Mean Square Deviation) and Fnat (Fraction of native contacts). However, RMSD-based measures may not be as accurate for cyclic peptides as Fnat measure[6], especially for peptides cyclized by disulfide bond, which structure may be partially incorrect at terminals, but despite that fact - at the main part of docked peptide it may still present the same interactions as interactions found in experimental structure. Due to that fact, in this work we used Fnat measure to verify docking success. Fnat was calculated as a fraction of experimental structure contacts reproduced in the generated model. The contact distance value for calculations in all-atom representation was defined as 8 Å. Generated models were classified according to CAPRI criteria: models with Fnat > 0.3 are considered acceptable, and models with Fnat > 0.5 are considered medium quality structures.

**Supplementary Results**

**Supplementary Table 1.** Success rate for docking peptides cyclized by their backbones to *holo* and *apo* receptor structures. The table reports Fnat values for the most accurate models: out from 10 top-scored models.

| PDB | Length | Global docking | | Local docking | |
|---|---|---|---|---|---|
| | | *holo* | *apo* | *holo* | *apo* |
| 1SFI | 14 | 0.14 | 0.11 | 0.37 | 0.28 |
| 3AV9 | 6 | 0.33 | 0.27 | 0.60 | 0.55 |
| 3AVA | 6 | 0.09 | 0.50 | 0.73 | 0.64 |
| 3AVB | 6 | 0.18 | 0.27 | 0.82 | 0.47 |
| 3AVF | 6 | 0.46 | 0.18 | 0.73 | 0.73 |
| 3AVG | 6 | 0.13 | 0.50 | 0.80 | 0.87 |
| 3AVH | 6 | 0.00 | 0.00 | 0.82 | 0.60 |
| 3AVI | 6 | 0.07 | 0.08 | 0.77 | 0.62 |
| 3AVJ | 6 | 0.21 | 0.07 | 0.64 | 0.43 |
| 3AVK | 6 | 0.64 | 0.14 | 0.71 | 0.50 |
| 3AVM | 6 | 0.82 | 0.19 | 0.73 | 0.46 |

| | | | | | |
|---|---|---|---|---|---|
| 3AVN | 6 | 0.50 | 0.10 | 0.90 | 0.50 |
| 3P8F | 14 | 0.25 | 0.07 | 0.44 | 0.40 |
| 3WNE | 8 | 0.63 | 0.64 | 1.00 | 1.00 |
| 3ZGC | 7 | 0.41 | 0.44 | 0.21 | 0.53 |
| 4K1E | 14 | 0.19 | 0.14 | 0.25 | 0.27 |
| 4KEL | 14 | 0.11 | 0.30 | 0.16 | 0.17 |
| 5XN3 | 8 | 0.21 | - | 0.32 | - |
| **Average** | | **0.30** | **0.24** | **0.61** | **0.53** |
| **Fnat > 0.3** | | **7** | **5** | **15** | **14** |
| **Fnat > 0.5** | | **4** | **3** | **12** | **10** |
| **Success rate** | | **39%** | **29%** | **83%** | **82%** |

**Supplementary Table 2.** Success rate for docking peptides cyclized by disulfide bond to *holo* and *apo* receptor structures. The table reports Fnat values for the most accurate models: out from 10 top-scored models.

| PDB | Length | Global docking | | Local docking | |
|---|---|---|---|---|---|
| | | *holo* | *apo* | *holo* | *apo* |
| 1HQQ | 13 | 0.30 | - | 0.80 | - |
| 1JBU | 15 | 0.07 | 0.04 | 0.30 | 0.38 |
| 1SMF | 9 | 0.20 | 0.27 | 0.16 | 0.38 |
| 1VPP | 19 | 0.50 | - | 0.50 | - |
| 2CK0 | 11 | 0.25 | - | 0.78 | - |
| 3G5V | 16 | 0.00 | - | 0.42 | - |
| 3P72 | 11 | 0.32 | 0.32 | 0.34 | 0.32 |
| 3WNF | 6 | 0.88 | 0.50 | 0.67 | 0.67 |
| 4IB5 | 13 | 0.04 | 0.50 | 0.76 | 0.55 |
| 4M1D | 14 | 0.57 | - | 0.18 | - |
| 4OU3 | 6 | 0.00 | 0 | 0.00 | 0.00 |
| 5CO5 | 16 | 0.25 | - | 0.31 | - |
| 5DJC | 13 | 0.18 | 0.22 | 0.22 | 0.20 |
| 5EOC | 13 | 0.14 | - | 0.58 | - |
| 5GRD | 10 | 0.23 | - | 0.40 | - |

| | | | | | |
|---|---|---|---|---|---|
| 5H5Q | 13 | 0.36 | 0.10 | 0.33 | 0.41 |
| 5TH2 | 12 | 0.25 | - | 0.45 | - |
| 5VB9 | 15 | 0.29 | - | 0.38 | - |
| 5WXR | 14 | 0.13 | 0.10 | 0.35 | 0.11 |
| 5XCO | 20 | 0.44 | 0.24 | 0.63 | 0.46 |
| **Average** | | **0.27** | **0.23** | **0.43** | **0.35** |
| **Fnat > 0.3** | | **6** | **3** | **16** | **7** |
| **Fnat > 0.5** | | **3** | **2** | **7** | **2** |
| **Success rate** | | **30%** | **30%** | **80%** | **70%** |

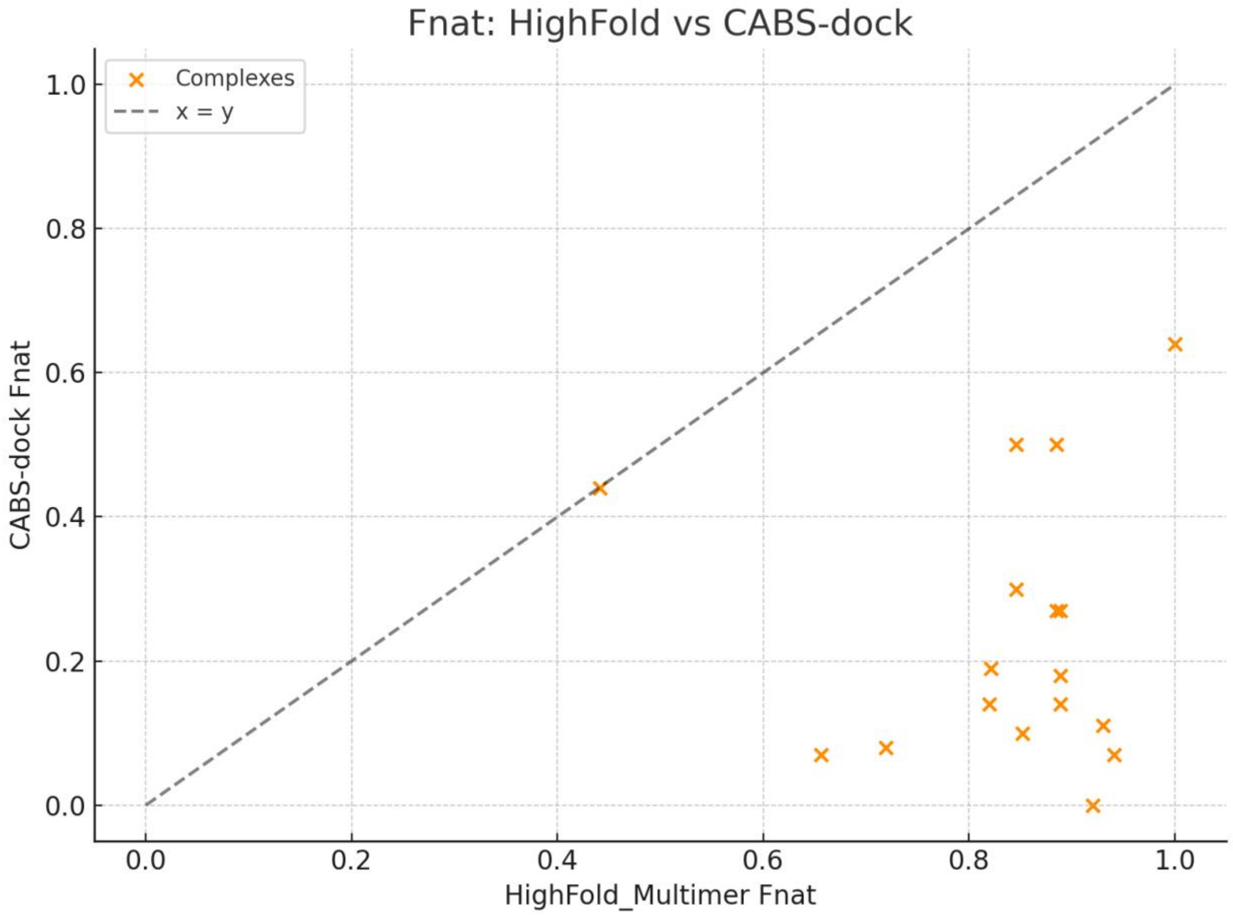


**Supplementary Figure 3.** Comparison of Top-1 Fnat values for CABS-dock and HighFold across 17 unbound cyclic protein-peptide complexes. Each point represents one complex, with the x-axis showing HighFold's Top-1 Fnat and the y-axis showing CABS-dock's Top-1 Fnat. The dashed identity line (x = y) indicates equal performance. The case of 3ZGC lies near the identity line (HighFold: 0.47, CABS-dock: 0.44), marking it as a particularly difficult target for both methods - yet one where CABS-dock reached much better quality in its Top-10 ensemble (Fnat up to 0.85 in the bound setup).
